# Brain functional connectivity initiates structured reorganization at a critical oxygen threshold during hypoxia

**DOI:** 10.1101/2025.08.18.670896

**Authors:** Daehun Kang, Koji Uchida, Clifton R. Haider, Norbert G. Campeau, Myung-Ho In, Erin M. Gray, Joshua D. Trzasko, Kirk M. Welker, Matt A. Bernstein, Max R. Trenerry, David R. Holmes, Michael J. Joyner, Timothy B. Curry, John Huston, Yunhong Shu

## Abstract

The human brain dynamically adapts to hypoxia, a reduction in oxygen essential for metabolism. The brain’s adaptive response to hypoxia, however, remains unclear. We investigated dynamic functional connectivity (FC) in healthy adults under acute hypoxia (FiO_2_ = 7.7%, 11.8%) using BOLD fMRI, physiological monitoring (PetO_2_, PetCO_2_, SpO_2_), and a Go/No-Go task. Principal component analysis identified a hypoxia-responsive FC component involving 400 cerebral parcels. This component emerged with a critical drop in PetO_2_ (∼53 mmHg), preceding changes in SpO_2_, BOLD signals, and behavior. These FC changes were network-specific and centered on the default mode network (DMN), which selectively synchronized with other high-level cognitive networks. In contrast, visual networks remained stable and segregated from the DMN. These results suggest that the brain proactively reorganizes its functional architecture in anticipation of oxygen decline, rather than in response to it. FC-based markers may offer early indicators of vulnerability in neurological or neurodegenerative conditions.

## 1 Introduction

As one of the most energy-demanding organs in the body, the human brain continuously anticipates and adapts to internal and external changes, including fluctuations in oxygen availability or hypoxia. Because oxygen is critical for oxidative metabolism, even a transient reduction can elicit a wide range of physiological and cognitive changes in the brain ^1, 2^.

Physiologically, a decline in arterial partial pressure of oxygen (PaO_2_) is the earliest and most sensitive indicator of hypoxic exposure ^3^. In response to reduced oxygen availability, the brain attempts to maintain metabolic homeostasis by increasing cerebral blood flow through vasodilation, along with adjustments in oxygen metabolism to sustain or prioritize energy production ^4, 5, 6, 7, 8^. Although compensatory, these physiological responses may not fully mitigate the effects of hypoxia, which can lead to temporary or permanent cognitive impairment and decline in neural function ^9, 10, 11, 12, 13, 14, 15^. These effects may emerge gradually or sequentially, and their onset and intensity often vary depending on the severity and duration of hypoxia exposure. Despite this, the neural mechanisms underlying the brain’s adaptive response to hypoxia, particularly during the early stages, remain poorly understood.

Functional connectivity (FC), defined as the temporal correlation of neural activity between brain regions, reflects the ongoing organization of the brain and has been widely used to characterize intrinsic neural organization and communication ^16, 17^. In the context of hypoxia, prior studies have increasingly explored how oxygen deprivation alters functional connectivity, using electroencephalography (EEG), functional near-infrared spectroscopy (fNIRS) or functional magnetic resonance imaging (fMRI) to demonstrate changes in FC after hypoxic exposure ^18, 19, 20, 21, 22, 23, 24^. EEG-based studies have reported region-specific changes in FC between frontocentral and occipitoparietal regions under hypoxic conditions. Acute hypoxia has been associated with increased frontocentral and reduced occipitoparietal integration in the theta band ^23^, whereas chronic hypoxia shows reduced frontal and increased occipital connectivity based on phase locking value, possibly reflecting long-term adaptation ^24^. fNIRS-based studies have suggested that the self-reference process remains robust under hypoxic conditions, reflecting the integrity of the default mode network (DMN) ^21^. Furthermore, research on patients with obstructive sleep apnea (OSA) reported subtle increases in fNIRS-based resting-state FC within the temporoparietal network, implying that repeated exposure to hypoxia may lead to functional reorganization ^22^.

fMRI-based studies of FC change after hypoxic exposure have demonstrated that even mild reductions in inspired oxygen levels can lead to significant reorganization of large-scale brain networks, for example, Liu et al. reported widespread network changes following acute exposure to 14.5% fractional inspired oxygen concentration (FiO_2_) ^18^. Similarly, long-term adaptation to high-altitude environments has been linked to increased FC within specific networks, such as the visual network ^19^. A preliminary analysis of the present study also revealed systematic, network-level reorganization during acute severe hypoxia exposure, with distinct patterns identified across major functional networks ^20^. These findings highlight the potential of fMRI, with its whole-brain coverage and high spatial resolution, to capture spatially distributed brain activity and dynamic functional connectivity during hypoxic challenges. Nonetheless, the brain’s real-time, network-level response to acute hypoxia remains poorly characterized. A more comprehensive understanding of these dynamic responses is necessary for understanding the brain’s vulnerability and adaptability under acute physiological challenges.

In studies of hypoxia-related brain responses, it is commonly interpreted that changes in FC occur consequent to reduction in oxygenated blood, often represented by peripheral oxygen saturation (SpO_2_). This interpretation is supported by evidence that reduced oxygen delivery to the brain directly alters synaptic connectivity ^25, 26^. Accordingly, FC changes have been traditionally considered secondary to physiological or structural alterations; a view consistent with current models of neuroplasticity ^27, 28^. However, the brain receives rapid and direct input through peripheral chemoreceptors, most notably the carotid bodies, which monitor PaO_2_ and signal through the glossopharyngeal nerve to the brainstem and higher centers involved in autonomic respiratory and cardiovascular regulation ^29, 30^. These pathways provide the brain with both subsequent downstream metabolic consequences and early direct sensory detection of metabolic challenge, i.e., hypoxia. Based on this, we propose an alternative hypothesis: FC changes may reflect a proactive neural reorganization in anticipating potential hypoxic threat, rather than merely a reactive response to it. In this study, we investigated the temporal dynamics and network-specific characteristics of FC in the human brain during acute hypoxic exposures. Specifically, we tested our hypothesis with particular attention to the relationship between FC reorganization and early physiological indicators such as end-tidal oxygen.

Direct measurement of PaO_2_ typically requires invasive arterial blood sampling, making real-time assessment impractical in neuroimaging studies. As a non-invasive proxy, the end-tidal partial pressure of oxygen (PetO_2_) offers a non-invasive and continuous estimate of systemic oxygen status ^31, 32^. Derived from exhaled air (EtO_2_), PetO_2_ correlates well with PaO_2_ under controlled conditions and captures early shifts in blood oxygenation. In this study, PetO_2_ served as a physiological marker to dynamically assess systemic oxygen status throughout the study.

To investigate the temporal dynamics of functional connectivity (FC) during hypoxia, we applied a causal sliding window approach to analyze the blood oxygenation level-dependent (BOLD) functional MRI data. This allowed us to track FC fluctuations as the subject underwent hypoxic challenge. The temporal association between the FC time series and the concurrently measured physiological and behavioral data were assessed to examine how brain network alterations respond to systemic and cognitive changes over time.

To further characterize temporal patterns of hypoxia-induced connectivity changes, we performed principal component analysis (PCA) on the time-resolved FC matrices. This data-driven approach allowed identification of the dominant spatiotemporal patterns of connectivity reorganization and their distribution across functional brain networks. To enable network-level analysis, standardized brain parcellation schemes based on large-scale population datasets were employed ^33, 34^. Together, these analyses provided a comprehensive view on how the brain dynamically adapts to oxygen deprivation in both time and space.

## 2 Methods and materials

### 2.1 Subjects

Eleven healthy subjects (six men, five women; 26.5 ± 4.5 years old) were recruited for this study ^8^. The study was conducted under an IRB-approved protocol, and all subjects provided informed written consent. All subjects were self-reported as healthy, with no known vascular, respiratory, cardiac or neurological disease. Prior to the experimental session, each subject was familiarized with the respiratory circuit and the cognitive task before they entered the MRI suite. Table 1 presents a summary of the physiological measures for the subject group.

**Table 1.** Physiological information for the subject group.

| Physiological measures | Mean ± SD |
| --- | --- |
| Height [cm] | 175 ± 10 |
| Weight [kg] | 81 ± 12 |
| Body mass index (BMI) [kg/m<sup>2</sup>] | 26 ± 3 |
| Systolic blood pressure (SBP) [mmHg] | 127 ± 14 |
| Diastolic blood pressure (DBP) [mmHg] | 79 ± 12 |
| Heart rate (HR) [bpm] | 70 ± 8 |

### 2.2 Experimental setup for hypoxia

To ensure subject safety, a board-certified anesthesiologist was present throughout all the MRI sessions, including hypoxic challenges, continuously monitoring the subjects’ cardiorespiratory parameters. Arterial blood pressure (BP) was monitored noninvasively using finger photoplethysmography (Nexfin, TPD Biomedical Instrumentation). Heart rate (HR), finger oxygen saturation (SpO_2_), breath-by-breath end-tidal O_2_ (EtO_2_) and CO_2_ (EtCO_2_) were continuously monitored throughout the study (Cardiocap/5, Datex-Ohmeda). Participants breathed through a low dead-space two-way non-rebreathing valve circuit for administration of gases and turbine system to measure tidal volume (Universal Ventilation Meter, Ventura, CA). Hypoxic gas was delivered from a gas reservoir (meteorological balloon) containing 7.7% or 11.8% oxygen, with the remaining balance consisting entirely of nitrogen, corresponding to severe and mild hypoxia conditions, respectively. During normoxia, 21% oxygen was administered instead.

Each participant underwent three 10-minute fMRI scans in the following order: normoxia, severe hypoxia, and mild hypoxia. During the normoxia and mild hypoxia runs, participants continuously received the assigned gas mixture throughout a 10-minute BOLD fMRI scan. In contrast, the severe hypoxia run followed a block design: participants first breathed normoxic air for 3 minutes (baseline), followed by 3 minutes of hypoxic gas administration, and then 4 minutes of re-oxygenation with normoxic air.

### 2.3 Data collections

#### MRI acquisition

All subjects were scanned on a compact 3T MRI scanner (C3T) equipped with a high-performance gradient system with 80 mT/m peak amplitude and 700 T/m/s peak slew rate ^35^. A 32-channel brain coil (Nova Medical, Inc. Wilmington, MA, USA) was used for brain imaging.

For functional scans, BOLD images were acquired using a gradient-echo echo-planar imaging (GRE-EPI) sequence over a 10-minute period. The imaging parameters were as follows: TR = 2 s, TE = 30 ms, isotropic spatial resolution = 2.5 mm, flip angle 90⁰, simultaneous-multi-slice (SMS) acceleration factor 3, no in-plane acceleration, and 300 repetitions.

For anatomical reference, a T1-weighted image (magnetization-prepared rapid acquisition with gradient echo, or MPRAGE) was obtained with the following parameters: TR = 5.4 ms, TE = 2.4 ms, inversion time TI = 1000 ms, flip angle (FA) = 8°, and isotropic spatial resolution = 1.0 mm. All images were retrospectively corrected for gradient nonlinearity distortions up to tenth order using the inline gradwarp algorithm ^36^.

#### Additional data collection

Independent measures of physiological signals, including EtO_2_, EtCO_2_, SpO_2_, cardiac pulsation, and respiration (via respiration belt), were continuously recorded during functional imaging.

### 2.4 Cognitive task

During the functional imaging sessions, the participants performed a computer-based Go/No-Go cognitive task developed by our team ^13, 14^. This cognitive test was developed using MATLAB (Matlab 9.5, R2019b, MathWorks, Natick, MA) software. The subjects were instructed to press a button in response to a “Go” stimulus but refrain from pressing the button for a “No-Go” stimulus.

Reaction time, commission error, and omission error were evaluated to assess behavioral performance during hypoxia. Reaction time was defined as duration from the appearance of a stimulus to the participant’s button press. Commission errors occurred when the participant incorrectly responded to No-Go stimuli, and omission errors when they failed to respond to Go stimuli. These metrics were continuously recorded throughout the cognitive test. To ensure participants were adequately familiarized with the task, a 10-minute practice session was conducted before the actual measurement, as recommended ^37^. Given our previous findings ^8, 13,14^, commission error rate was selected as the primary behavioral marker of hypoxia-induced cognitive decline.

### 2.5 Data Processing

#### Preprocessing for functional images

Prior to analyzing the dynamic FC with the sliding window, the functional data with 300 consecutive volumes of EPI images were preprocessed. The preprocessing steps included initial 10-volume truncation, de-spiking, slice-timing correction, physiological artifact correction (RETROICOR) ^38, 39^, EPI-MPRAGE alignment, inter-EPI-volume registration, spatial smoothing by a Gaussian kernel with a 4 mm full-width-at-half-maximum (FWHM) on volume datasets, and signal scaling to 100. Next, Legendre polynomial (up to 4 order), motion, white matter/cerebral spinal fluid (WM/CSF) (ANATICOR and CompCor) linear regressors ^40, 41^, were applied to obtain residual BOLD signal using the AFNI software package ^42^. All EPI data were evaluated for abrupt head motion artifacts, passing the sudden motion and outliner detections of AFNI at the threshold level of 0.3 and outlier fraction of 0.1 for the Euclidean L2 norm of motion displacement during each TR, respectively.

#### Processing for anatomy image

The T1-weighted anatomical images were preprocessed to segment brain structures using FreeSurfer software ^43^, which were then used for registration to the EPI images and to the standard cerebral surfaces provided by AFNI and SUMA software ^42^.

#### Pre-defined ROI with intrinsic brain networks

Cerebral regions of interest (ROI) were defined based on the 400 cerebral parcellations suggested by Schaefer’s atlas ^34^, available on the AFNI standard surface model (https://afni.nimh.nih.gov/pub/dist/atlases/SchaeferYeo/). The 400 cerebral patches on the surface space were transferred onto the individual image volumetric space for EPI datasets. Each ROI was associated with one of 17 intrinsic connectivity networks, enabling both ROI-level and network-level analysis of functional connectivity. Connectivity measures were assessed both within (intra-) and between (inter-) networks.

#### Dynamic changes by sliding window

To examine dynamic FC, average time courses for each of the 400 ROIs were extracted from the preprocessed residual BOLD signals. FC between ROIs were calculated by Fisher-z-transformed Pearson correlation using MATLAB software. The time-resolved connectivity information was derived using a 90-second causal sliding window approach and a step size of a 2-second shift, yielding 246 overlapping time windows per scan. For each window, a full ROI-by-ROI FC matrix was computed for all 400 cortical ROIs. In total, 79,800 unique ROI-to-ROI connections were computed per window. This process generated a time series of dynamic FC matrices for each subject.

#### Principal component analysis

To identify temporal connectivity patterns associated with hypoxia, PCA was only applied to the fMRI session that included a block-design paradigm (normoxia–severe hypoxia–normoxia). This session was chosen due to its well-defined temporal structure, which facilitates the extraction of hypoxia-related patterns. PCA was conducted with MATLAB (R2019b) with default settings. The input for PCA consisted of a matrix of 2,706 time points × 79,800 functional connections. The time points were derived from 246 sliding windows across 11 subjects and concatenated together. In this framework, each time point represented an observation, and each FC was treated as a variable. Principal components were ranked by their explained variance, and dominant components were selected for further analysis. These resulting components were then reshaped into a 246 (time points) × 11 (subjects) matrix to allow for inspection of their temporal dynamics across individuals.

#### Monte-Carlo permutation test

Coefficients of the most dominant principal component across 79,800 functional connections were ranked by absolute value, and the top 5% (3,990 edges) were identified as hypoxia-responsive. These edges were mapped onto a 17 × 17 network matrix (Schaefer atlas) to quantify their distribution across network pairs. Statistical significance was assessed using a one-tailed Monte Carlo permutation test (10,000 iterations). In each iteration, 3,990 edges were randomly sampled from the 79,800 using MATLAB’s *randperm* (R2019b) and assigned to the network matrix. P-values were computed as the proportion of permuted counts greater than or equal to the observed value, using the standard formula: (k + 1)/(N + 1), and were corrected for multiple comparisons across 153 cells using the Benjamini– Hochberg false discovery rate (MATLAB’s *mafdr*). The same procedure was applied to the bottom 10% (stable FC edges) and other percentile thresholds to assess their statistical significance.

## 3 Results

### 3.1 Effects of acute severe hypoxia on multiple measures

Fig. 1 provides the average time courses across subjects, illustrating the physiological, behavioral, and neural responses to acute severe hypoxia. Fig. 1a and 1b depict the experimental paradigm and the timing of hypoxia stimulation, respectively. Fig. 1(c-f) provides physiological responses, including time courses of PetO_2_, PetCO_2_, peripheral SpO_2_, and the bulk BOLD signal, respectively. Fig. 1g displays changes in behavioral performance, by representing the change in commission error rate, while Fig. 1h provides the corresponding dynamics of mean FC. Each time point in Fig. 1g and 1h was computed using a 90-second causal sliding window, either by averaging (Fig. 1g) or by calculating Fisher-Z transformed Pearson correlation coefficients (Fig. 1h). The time series represent group-level averages across all subjects (N = 11).

**Fig. 1.**
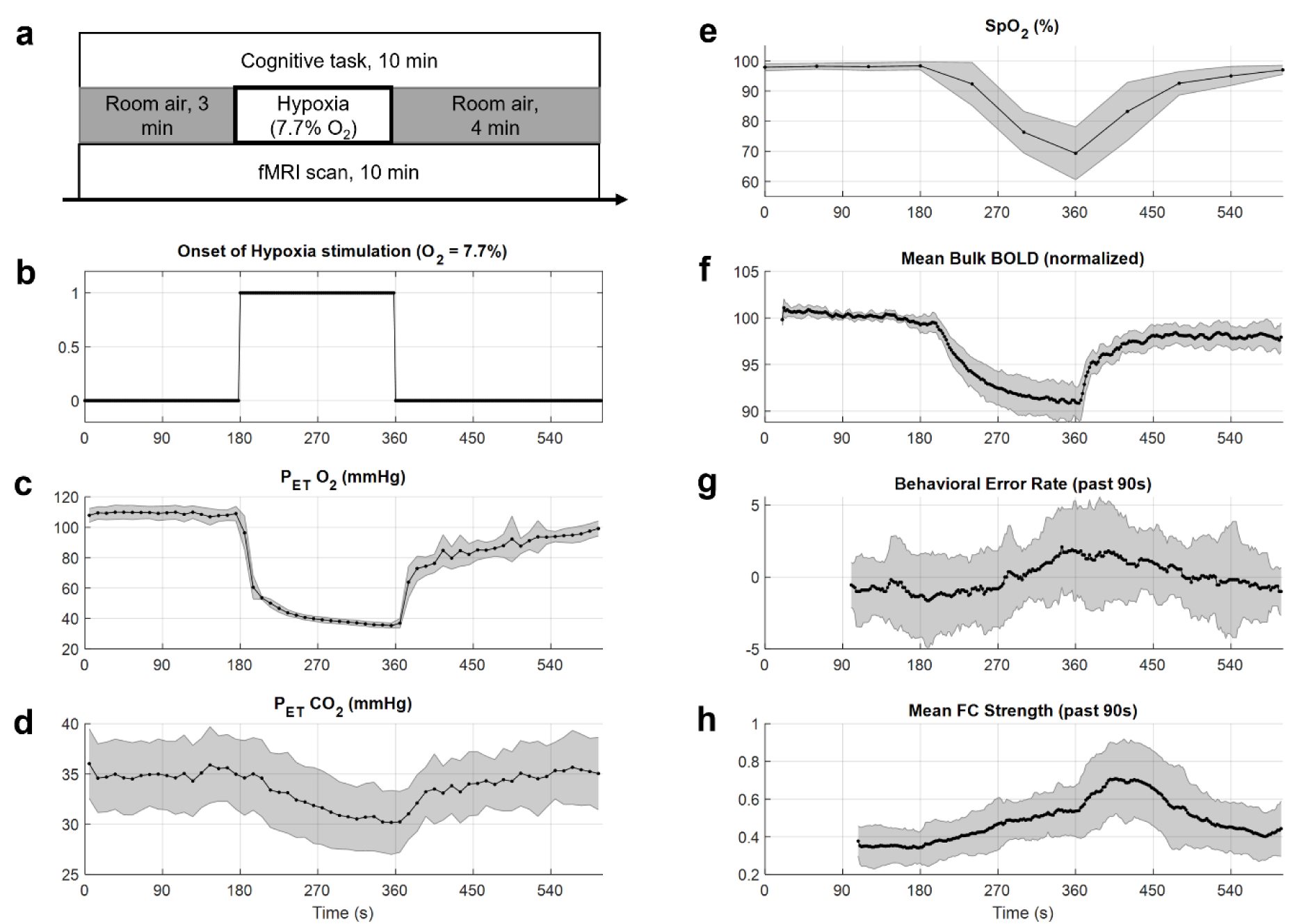
Multi-domain effects of acute severe hypoxia on respiratory gas dynamics, oxygenation responses, cognition performance, and brain connectivity. **a**, Experimental protocol with a 10-minute BOLD fMRI session including 3 minutes of normoxia, 3 minutes of hypoxia (FiO_2_ = 7.7%), and 4 minutes of normoxia reoxygenation, performed concurrently with a Go/No-Go cognitive task. **b**, Time course marking the onset and duration of hypoxic stimulation. **c, d**, Time course of estimated partial pressures of end-tidal oxygen (PetO_2_) and carbon dioxide (PetCO_2_), respectively. **e**, Time course of peripheral oxygen saturation (SpO_2_). **f**, Hypoxia-induced bulk BOLD signal normalized to the 1-minute average proceeding hypoxia onset. **g**, Behavioral change, expressed in the change in commission error rate (Δcommission error) over a 90-second causal sliding window. **h**, Temporal dynamics of the global functional connectivity within a 90-second causal sliding window. Sampling intervals: 2 seconds for the panels (a, f, g, h); 10 seconds for (c, d); 30 seconds for (e). Error bar (grey area) denotes standard deviation across subjects (N = 11).

PetO_2_ exhibited the most rapid response to severe hypoxia, dropping sharply from 96.3 to 53.7 mmHg within 20 seconds of onset followed by a continued exponential-like decline. In contrast, PetCO_2_ decreased gradually and approximately linearly throughout the hypoxic period. Although both measures began to recover immediately after hypoxia ended, neither fully returned to baseline within the subsequent 4 minutes of re-oxygenation period.

Compared to PetO_2_, peripheral SpO_2_ declined more gradually and began a slow recovery after the hypoxia period. However, due to its relatively coarse sampling interval (30 seconds), precise temporal comparisons with other physiological signals were limited. In contrast, the mean bulk BOLD signal exhibited a delayed decline in an exponential fashion, with an onset lag of approximately 22 seconds after hypoxia onset. During re-oxygenation, the bulk BOLD signal increased sharply, also in an exponential fashion, following a brief delay of about 8 seconds.

Behavioral performance during the Go/No-go task, to assess cognitive deterioration, was evaluated using the commission error rate ^8, 13, 14^. The relative error rate began to increase approximately 90 seconds after hypoxia onset and gradually decreased after the hypoxia period ended.

Global mean dynamic FC strength increased shortly after hypoxia onset and began to decrease around 40 seconds after the hypoxic stimulus ended. Compared to the bulk BOLD signal and the behavioral performance, FC changes occurred earlier and exhibited the slowest recovery (i.e., delayed decrease in FC strength), indicating a prolonged neural response to hypoxic stress.

### 3.2 Hypoxia-responsive changes in functional connectivity

To analyze temporal change in FC induced by acute severe hypoxia, PCA was performed to the time-resolved FC matrices. As outlined in Fig. 2a, the table summarizes the top five principal components with the highest explained variance. The first principal component (PC1) alone accounted for 38.9% of the total variance, whereas each of the remaining components contributed less than 4.4%, indicating the dominance of PC1. Fig. 2b plots the time course of PC1, with a red horizontal bar indicating the hypoxia period. PC1 time course exhibited a marked increase shortly after the severe hypoxia onset, suggesting that PC1 captures the principal mode of FC reorganization driven by the hypoxia.

**Fig. 2.**
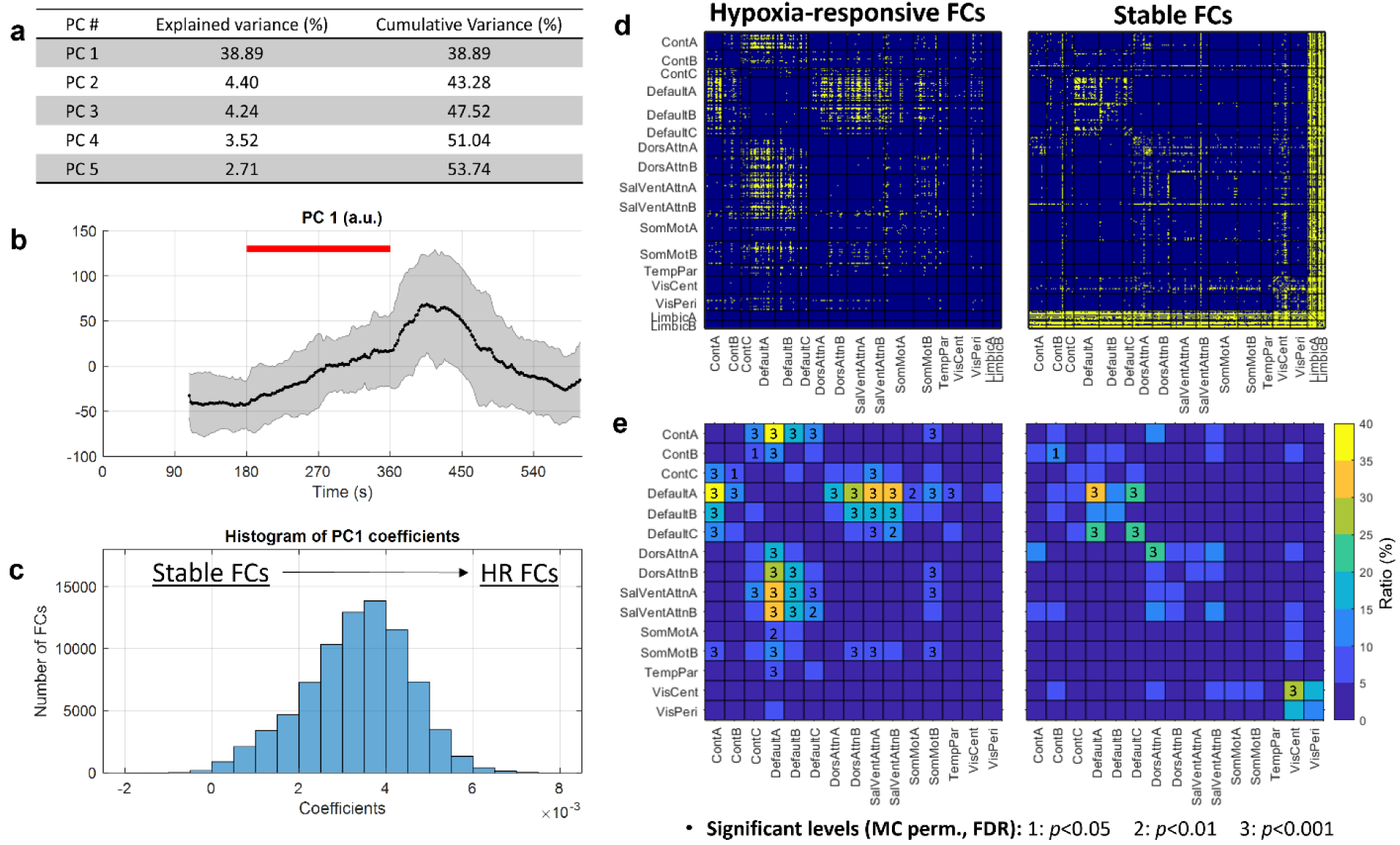
Hypoxia-responsive changes in functional connectivity are network-specific. To examine hypoxia-induced temporal dynamics in brain functional connectivity, principal component analysis (PCA) was applied to dynamic connectivity profiles comprising 79,800 pairwise connections among 400 cortical parcels (Schaefer atlas) across subjects. **a**, Summary table presenting the explained and cumulative variance of the top five principal components (PCs), with PC1 accounting for the largest portion of the variance. **b**, Group-averaged time series of PC1. The red horizontal bar indicates the period of acute severe hypoxia. Shaded areas denote standard deviation across subjects. The y-axis represents signal intensity in arbitrary units (a.u.). **c**, Histogram of PC1 coefficients. For further analysis, functional connections (FCs) with the top 5% of largest coefficients were classified as hypoxia-responsive (HR) FCs, and the bottom 10% with the smallest coefficients were defined as stable FCs. **d**, 400 × 400-ROI FC matrices, visualizing the distribution of HR FCs (left) and stable FCs (right). **e**, Inter- and intra-network distributions of HR and stable FCs. Cell colors indicate the percentage of possible connections within and between the 17 canonical brain networks. Numbers shown in cells represent significance levels from Monte Carlo (MC) permutation tests (1: p<0.05, 2: p<0.01, 3: p<0.001), corrected for multiple comparisons using the Benjamini–Hochberg false discovery rate (FDR) method. Notably, the Default A network exhibited selective increases in inter-network connectivity, with minimal change in intra-network connectivity. Limbic A and B networks were excluded from this summary due to signal dropout in many of their parcels. Abbreviations of brain networks: Cont = Control; Default = Default Mode; DorsAttn = Dorsal Attention; Limbic = Limbic; SalVentAttn = Salience/Ventral Attention; SomMot = Somatomotor; TempPar = Temporoparietal; VisCent = Visual Central; VisPeri = Visual Peripheral.

Fig. 2c highlights the distribution of the PC1 coefficients across all 79,800 connections. The histogram approximated a unimodal (near Gaussian) distribution centered at 3.3 ± 1.2 (×10^-3^), with a minor negative tail of −0.2 ± 0.2 (×10^-3^), which was considered negligible. FC edges with strongly positive PC1 coefficients were classified as hypoxia-responsive (HR-FCs), while those with near-zero coefficients were considered stable (stable-FCs). Given the absence of a natural cutoff in the distribution, the top 5% and bottom 10% of FC edges were operationally selected based on the ranked absolute PC1 coefficients, yielding 3,990 HR-FCs and 7,980 stable-FCs out of all possible 79,800 connections. Fig. 2d visualizes the subset HR-FCs and stable-FCs as yellow dots in 400 × 400 FC matrices. FCs were grouped by the 17 intrinsic brain networks from the Schaefer atlas to reveal spatial and network-specific patterns.

Most HR-FCs were predominantly observed in inter-network connections involving the Default A and B networks and their connections to Control A and B, Dorsal Attention A and B, and Salience/Ventral Attention A and B networks. In contrast, stable-FCs were largely confined to connections associated with Limbic A and B networks, as well as some intra-network connections within Default A and other networks.

Fig. 2e provides a network-level summary of Fig. 2d, where the colors represented the proportion of HR-FCs and stable-FCs relative to the total number of possible FC edges within or between networks. ROIs belonging to the Limbic A and B networks, as defined in Schaefer’s atlas, were excluded from this panel due to severe signal dropout in the susceptibility-prone regions of these networks such as orbitofrontal cortex and temporal pole (labeled by OFC_x and TempPole_x, respectively, in the Schaefer’s atlas).

The numbers provided in each cell of Fig. 2e indicate significance levels obtained from Monte Carlo permutation tests: 1 indicates *p* < 0.05, 2 indicates *p* < 0.01, and 3 indicates *p* < 0.001 (Benjamini–Hochberg false discovery rate-corrected). Based on the permutation results, Default A network exhibited the most extensive hypoxia-related increases in inter-network connectivity, while its intra-network connectivity remained largely preserved, consistent with its role as a stable structural and functional core. These findings suggest that Default A network may serve as a central coordinator during the systemic hypoxic perturbation. Notably, while the Default A network exhibited increased connectivity with most other networks, it exhibited little to no significant hypoxia-responsive interaction with the Control C, Visual Central, and Visual Peripheral networks. This selective pattern suggests a targeted reorganization rather than a global connectivity shift.

### 3.3 A critical oxygen threshold triggers abrupt changes in functional connectivity

Fig. 3(a, c, f) illustrates the experimental paradigms for normoxia, acute severe hypoxia, and mild hypoxia, respectively. A block design paradigm was employed only in the acute severe hypoxia condition, whereas normoxia and mild hypoxia (FiO_2_ = 21.0% and 11.8%, respectively) were assessed under continuous exposure without block structuring.

**Figure 3.**
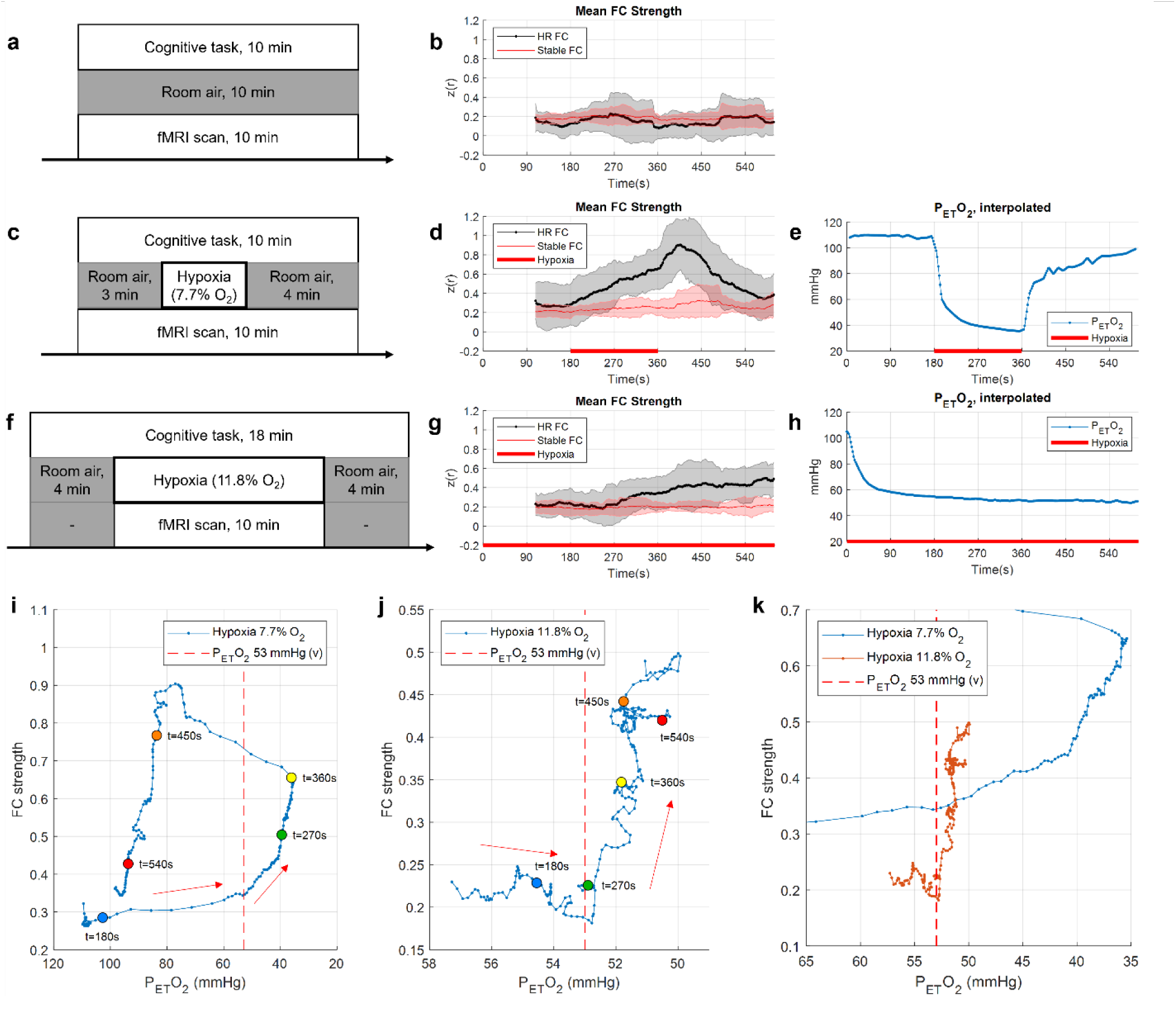
Hypoxia-induced changes in functional connectivity are triggered when PetO_2_ crosses a critical threshold. Normoxia and mild hypoxia experiments were included to validate the hypoxia-responsive (HR) change in FC. **a, b**, fMRI protocol under normoxia (FiO_2_ = 21.0%) and the corresponding dynamics of hypoxia-responsive (HR) and stable FCs. **c-e**, fMRI protocol under severe hypoxia (FiO_2_ = 7.7%) and the corresponding dynamics of HR and stable FCs and interpolated PetO2 time course. **f-h**, fMRI protocol under mild hypoxia (FiO_2_ = 11.8%) and the corresponding dynamics of HR and stable FCs and interpolated PetO2 time course. **i, j**, Scatterplots of mean HR-FC strength versus PetO_2_ during severe and mild hypoxia, respectively. Colored dots (blue to red) indicate five time points from 180s to 540s (90-second intervals), illustrating temporal progression alongside red arrows. In panel **i**, the blue and yellow dots mark the onset and offset of severe hypoxia, respectively. In panel **j**, all dots represent time points under hypoxia, as mild hypoxia was applied throughout. A vertical dashed line at 53 mmHg indicates the proposed threshold at which HR-FCs begin to emerge. **k**, Combined view of i and j, showing the consistent threshold effect under both hypoxia conditions, suggesting a common PetO_2_ tipping point for triggering FC reorganization.

Based on the HR-FCs and stable FCs previously identified under severe hypoxia, we examined these FC responses under all three hypoxia conditions. Fig. 3b illustrates the temporal dynamics of the two FC edge groups during normoxia, while Fig. 3d and 3g present those from the severe and mild hypoxia conditions, respectively. HR-FCs under normoxia exhibited no noticeable changes over time, whereas HR-FCs under severe hypoxia increased sharply following the onset of hypoxia, in alignment with the PC1. Interestingly, under mild hypoxia, although the reduced oxygen level was applied continuously, HR-FCs began to increase approximately 276 seconds after onset. Across all three conditions, stable FCs demonstrated no appreciable changes over time.

Fig. 3e and 3h plot the PetO_2_ time courses for the severe and mild hypoxia conditions, respectively, linearly interpolated to match the temporal resolution of FC dynamics. These PetO_2_ curves were projected on top of the corresponding dynamic FC responses in Fig. 3d and 3g, and the plots are provided in Fig. 3i and 3j, respectively. In severe hypoxia condition (Fig. 3i), PetO_2_ dropped rapidly right after hypoxia onset, and HR-FC strength began to rise sharply when PetO_2_ fell below approximately 53 mmHg. Colored dots (blue to red) indicate five time points from 180s to 540s (90-second intervals), illustrating temporal progression alongside red arrows. The green and yellow dots mark the onset and offset of severe hypoxia. A similar threshold-driven response was also observed under mild hypoxia (Fig. 3j), where HR-FCs began to increase once PetO_2_ dropped below approximately 53 mmHg. While the same five color-coded time points are provided in Fig. 3j, there was no distinct onset or offset under the mild hypoxia condition. Fig. 3k combines these results from Fig. 3i and 3j, demonstrating that in both paradigms, HR-FC changes consistently began around a critical threshold of PetO_2_ = 53 mmHg.

## 4 Discussion

In this study, we examined how acute hypoxia influences large-scale brain FC using fMRI under normoxia, mild hypoxia, and severe hypoxia conditions. Dynamic FC analysis and PCA revealed that 3-minute severe hypoxia induced network-specific reorganization, particularly centered on the DMN (Default A), as shown in Fig. 2. These FC changes emerged prior to measurable declines in SpO_2_, bulk BOLD signal, or behavioral performance as highlighted in Fig. 1. Among the monitored physiological markers, end-tidal oxygen pressure (PetO_2_) exhibited a distinct threshold-triggered association with FC changes during both severe and mild hypoxia (Fig. 3i and 3j). While hypoxia-responsive FC edges might superficially resemble reactive responses, the concurrent presence of stable FC edges, particularly those consistently preserved throughout the hypoxic period, suggests a proactive and structured reconfiguration of network connectivity in response to impending oxygen stress.

The most notable FC-related finding is that hypoxia-responsive changes were not uniformly distributed across the large-scale brain networks but instead show distinct network-level specificity. Specifically, strong FC increases were observed between Default A and other high-level networks such as Control A/B, Salience/Ventral Attention A/B, and Dorsal Attention A/B networks, etc. This organization suggests an expansion of Default A’s role, not through global connectivity increases, but via selective integration with high-level cognitive control and attention networks. This pattern was also consistently observed across multiple coefficient thresholds (Supplementary Fig. 1), further reinforcing the robust and central role of Default A in hypoxia-related network reorganization. This may resonate with prior findings that self-referential processing remains robust under hypoxia, as observed in an fNIRS study ^21^, possibly reflecting the preserved function of DMN-related networks.

In distinction, Default A exhibited relatively fewer hypoxia-responsive FC edges with networks such as Control C, and Visual Central/Peripheral networks. Additionally, stable FCs, those less affected by hypoxia, were primarily observed within the intra-network connections of the Default A network. The Control C network includes parcels located in the precuneus and posterior cingulate cortex (labeled *pCun_* and *Cingp_*, respectively), which anatomically overlap with key regions of the Default A network ^34^. This overlap may indicate functional integration with Default A, potentially rendering these regions less responsive to further reorganization under hypoxia.

Visual Central/Peripheral networks were among the few networks not included in the Default A-driven selective integration during hypoxia. Interestingly, this observation aligns with EEG studies reporting hypoxia-induced increases in functional integration within the θ band, particularly in frontocentral regions, accompanied by decreased integration in occipitoparietal regions ^23^. Functional segregation between frontal and occipital regions has also been observed in long-term hypoxia studies, both in humans exposed to high-altitude conditions ^24^ and in animal models under experimentally induced hypoxia ^44^. While EEG studies have reported increased FC within occipital regions under hypoxia, particularly in the β band or when measured using phase locking value, our fMRI data demonstrated a high proportion of stable FCs within the visual networks. In particular, the Visual Central network remained one of the most functionally segregated networks, consistently showing low involvement across all coefficient thresholds (see Supplementary Fig. 1). This discrepancy may be attributed to EEG-based FC reflecting frequency-specific, local neuronal processes that are not captured by BOLD-based FC measures used in this study.

Although both PetO_2_ and PetCO_2_ decreased during hypoxia, only PetO_2_ demonstrated a clear threshold-related association with FC changes. Since CO_2_ is a potent vasodilator that influences the BOLD signal, FC changes may be confounded by declining PetCO_2_ during hypoxia. ^45, 46^. However, prior work indicates that WM/CSF regression can approximate the effect of PetCO_2_ clamping ^46^. Our preprocessing pipeline for functional MRI datasets incorporated ANATICOR ^40^ and CompCor ^41^, which likely mitigated CO_2_-related confounds.

Global signal regression was not applied. As provided in Supplementary Figure 2, although some increases in FC strength coincided with declining PetCO_2_ during severe hypoxia, the overall relationship was weak and inconsistent. Under mild hypoxia, PetCO_2_ fluctuations were minimal and demonstrated no discernible correspondence with FC changes. These findings suggest that PetCO_2_ did not play a primary role in driving FC reorganization. Nonetheless, since our data captured only a decreasing trend in PetCO_2_, further studies are required to characterize the brain’s network-level response to elevated CO_2_ levels ^47^. Taken together, our results suggest that proactive FC reorganization was more closely associated with changes in PetO_2_ than in PetCO_2._ Further work is needed to disentangle the specific contributions of O_2_ and CO_2_ to hypoxia-induced network dynamics.

The brain’s prioritization of O_2_ monitoring at the network level may stem from the fact that O_2_ is an essential external resource that cannot be synthesized or stored, and its deficiency poses an immediate threat to neuronal viability and synaptic function ^22, 25, 26^. In contrast, CO_2_ is an internally generated metabolic byproduct that can be buffered and expelled. Although it has long been a central focus of neurovascular regulation due to its strong vasodilatory effects and role in pH modulation, it does not represent a resource whose absence directly threatens cellular function. From this perspective, O_2_ serves as the brain’s metabolic budget ^1^: any decrease in its availability must be detected early and acted upon decisively. Thus, the observed anticipatory reorganization at the large-scale network level, beyond localized or reactive cellular responses, may reflect an intrinsic preparedness to mitigate impending metabolic stress.

Interestingly, hypoxia-induced FC change did not return to baseline immediately, even after PetO_2_, SpO_2_, and bulk BOLD signals had largely recovered from severe hypoxia. Notably, the fMRI study also reported no correlation between altered FC and SpO_2_ following hypoxia ^18^. These findings suggest that, although the hypoxia-induced FC responses are initiated proactively in response to a critical drop in PetO_2_, their evolution and recovery may follow a distinct regulatory trajectory that is decoupled from the restoration of other physiological signals, including PetO_2_ itself. However, since our observations were derived exclusively from a single severe hypoxia condition, future studies involving varying levels or types of hypoxic exposure are needed to clarify the robustness and specificity of the observed FC recovery process.

The proactive framework may offer a novel avenue for evaluating physiological or therapeutic interventions. Specifically, if the brain exhibits proactive network-level adaptations in response to external changes, then it may be possible to assess treatment efficacy before overt behavioral changes occur. Such early neural signatures could reduce reliance on delayed and risk-bearing behavioral endpoints and provide a more sensitive, system-level biomarker. If proactive reorganization reflects a general systems-level strategy, it could serve as an early indicator of therapeutic engagement, even in the absence of immediate behavioral changes FC-based metrics may thus offer a valuable window into brain-level responsiveness, enabling earlier and potentially safer assessments of treatment effects.

The findings suggest that FC reorganization in response to acute physiological stress may reflect a broader systems-level principle, wherein the brain leverages proactive, structured adaptations to preserve critical functions amid external or internal disruption. A comparable process may underlie early pathological changes in Alzheimer’s disease ^22, 23^, as proposed in the cascading network failure model (Jones, 2016), which describes how functional decline in the posterior DMN (pDMN) triggers increased synchronization with other DMN subsystems, including the ventral (vDMN) and anterior-dorsal (adDMN) components. This reorganization in turn imposes an excessive burden on the hippocampus, potentially accelerating pathological progression. While traditionally viewed as a consequence of pathology, such network-level restructuring may reflect a proactive, but ultimately maladaptive, response to emerging neural dysfunction. In contrast to hypoxia, where FC changes are transient and potentially reversible, the changes in Alzheimer’s appear to become progressively entrenched, as they are typically observed after clinical symptoms have already emerged. Nonetheless, interpreting these processes through a proactive adaptation framework may help reveal early vulnerability in neural systems, potentially before irreversible dysfunction becomes apparent. This perspective may inform the use of FC-based imaging biomarkers or neurostimulation techniques (e.g., transcranial magnetic stimulation ^48^) for preclinical detection and therapeutic targeting in neurodegenerative disorders.

Despite these insights gained from this study, several limitations should be acknowledged. First, the sample size was relatively small. While dense window shifts (2-second increments) and PCA centered on PC1 allowed robust group-level dynamic FC pattern detection, larger cohorts are needed to assess the generalizability, especially for additional principal components beyond PC1. Second, FC was estimated during a Go/No-Go task, in which short-duration stimulus events were presented at fixed pseudo-random intervals across sessions, ensuring consistent task structure. While this design controlled for task-related effects, it remains possible that different cognitive tasks or resting-state paradigms could yield distinct baseline FC patterns and modulate the brain’s response to hypoxia. Nevertheless, the consistency of primary component (PC1) across conditions suggests a dominant contribution from the hypoxia itself, meriting further investigation. Finally, while we observed a selective increase in inter-network connectivity centered on the Default A network during hypoxia, the functional significance of this reorganization remains unclear. Given that FC reflects statistical associations rather than direct neural communication, future studies employing complementary modalities, such as metabolic imaging/assessment or electrophysiology, will be essential to clarify the underlying mechanisms and functional implications of this network-level reconfiguration.

Acute hypoxia presents a severe external challenge to the brain’s oxygenation and metabolic stability. Our findings demonstrate that the brain not only reacts to falling oxygen levels, but also engages in a structured, proactive reorganization of FC. Centered on the DMN, this adaptation is triggered by a critical PetO_2_ threshold and precedes changes in other physiological and behavioral markers. The delayed recovery of FC, relative to other physiological signals, underscores the distinct regulatory trajectory of network-level dynamics. Understanding such proactive adaptations may offer new insights into the principles of neural resilience and vulnerability across both healthy and pathological states.

## 5 Ethics statement

The study was approved by the Institutional Review Board of the Mayo Clinic and written informed consent was obtained from all participants.

## Acknowledgements

The authors acknowledge the contributions of the Human Integrative Physiology Laboratory, Clinical Research and Trials Unit and Special Purpose Processor Development Group at the Mayo Clinic. This work was supported by the Office of Naval Research and by National Institute of Biomedical Imaging and Bioengineering (NIBIB) at the National Institutes of Health (NIH). The opinions, findings and conclusions or recommendations expressed in this material are those of the author(s) and do not necessarily reflect the views of the Office of Naval Research.

## 6 Author contributions

**Daehun Kang**: Conceptualization, Methodology, Formal analysis, Writing - Original Draft, Visualization, **Koji Uchida**: Conceptualization, Methodology, **Clifton R. Haider**: Conceptualization, Methodology, Project administration, Funding acquisition, **Norbert G. Campeau:** Conceptualization, Methodology, **MyungHo In:** Methodology, **Erin M. Gray:** Investigation, **Joshua D. Trzasko:** Methodology, **Kirk M. Welker:** Methodology, **Matt A. Bernstein:** Methodology, Supervision, **Max R. Trenerry:** Conceptualization, Methodology, **David R. Holmes:** Conceptualization, Methodology, **Michael J. Joyner:** Conceptualization, Methodology, **Timothy B. Curry:** Conceptualization, Methodology, Project administration, Funding acquisition, Supervision, **John Huston III:** Conceptualization, Project administration, Supervision**, Yunhong Shu:** Methodology, Writing - Original Draft, **All**: Writing - Review & Editing

## 9 Data availability statements

The datasets generated and analyzed in this study are not publicly available due to institutional regulations and data protection policies governing protected health information.

## 10 Declaration of generative AI and AI-assisted technologies in the writing process

During the preparation of this work the author used ChatGPT / GPT-4o in order to improve language and readability. After using the ChatGPT / GPT-4o, the authors reviewed and edited the content as needed and take full responsibility for the content of the publication.

