## Supplemental Figures for "Brain functional connectivity initiates structured reorganization at a critical oxygen threshold during hypoxia"

### Supplementary materials

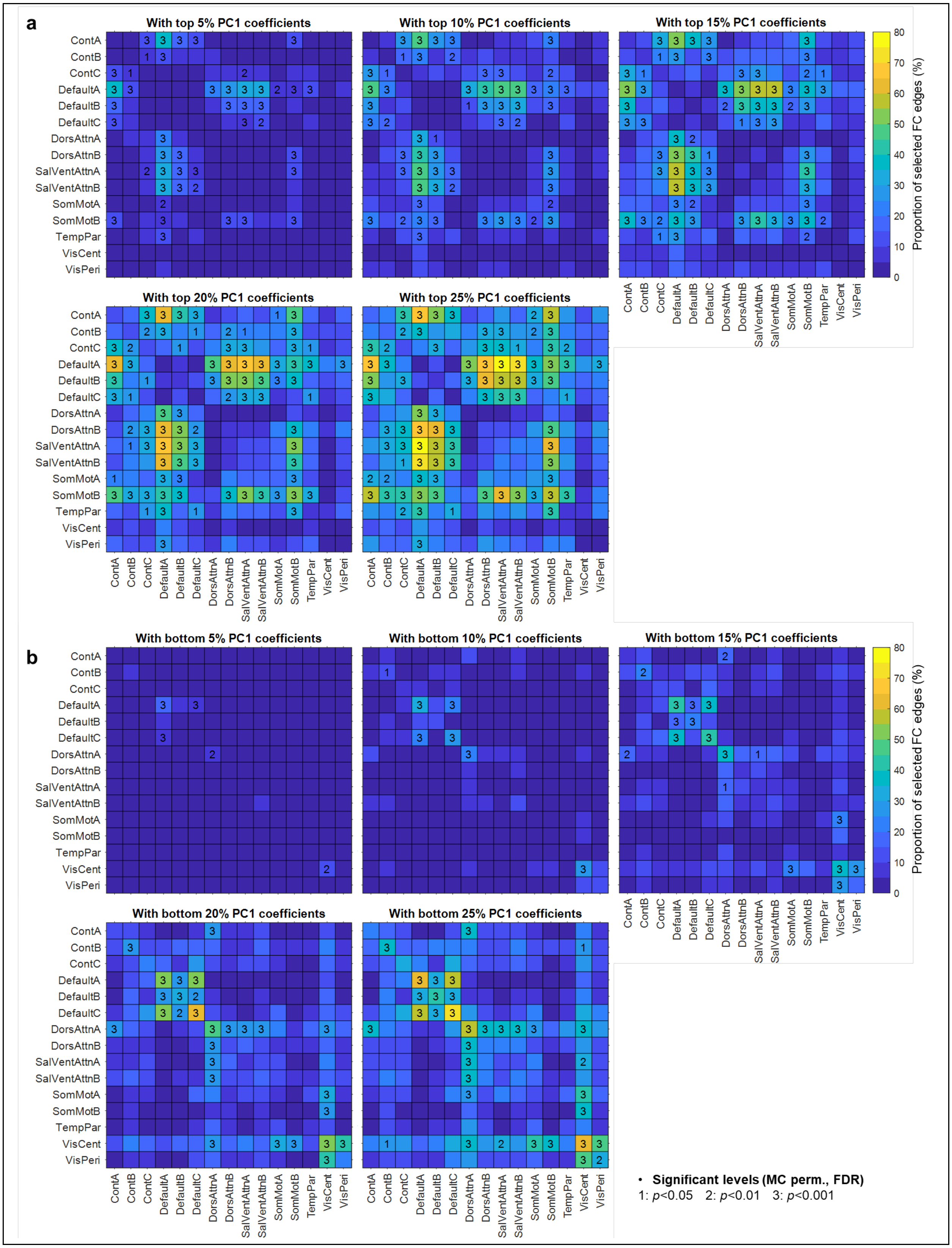

**Supplementary Figure 1. Proportion of functional connectivity edges selected from top and bottom PC1 coefficients across multiple thresholds.**

The top (a) and bottom (b) percentiles of PC1 coefficients (as shown in Fig. 2c) were used to statistically identify the hypoxia-responsive and stable network-wise functional connections, respectively.

Each matrix depicts the proportion of selected FC edges within and between the 17 canonical brain networks, evaluated across five coefficient thresholds: 5%, 10%, 15%, 20%, and 25%.

Cell colors indicate the relative proportion of selected edges, while numbers shown in cells represent significance levels from Monte Carlo (MC) permutation tests (1: $p<0.05$, 2: $p<0.01$, 3: $p<0.001$), corrected for multiple comparisons using the Benjamini–Hochberg false discovery rate (FDR) method.

Default A consistently exhibited a high proportion of selected FC edges across thresholds, indicating its dominant role in hypoxia-related network reorganization. In contrast, visual networks, especially VisCent, remained functionally segregated with minimal involvement across all conditions.

Limbic A and B networks were excluded from this summary due to signal dropout in many of their parcels.

Abbreviations of brain networks: Cont = Control; Default = Default Mode; DorsAttn = Dorsal Attention; Limbic = Limbic; SalVentAttn = Salience/Ventral Attention; SomMot = Somatomotor; TempPar = Temporoparietal; VisCent = Visual Central; VisPeri = Visual Peripheral.

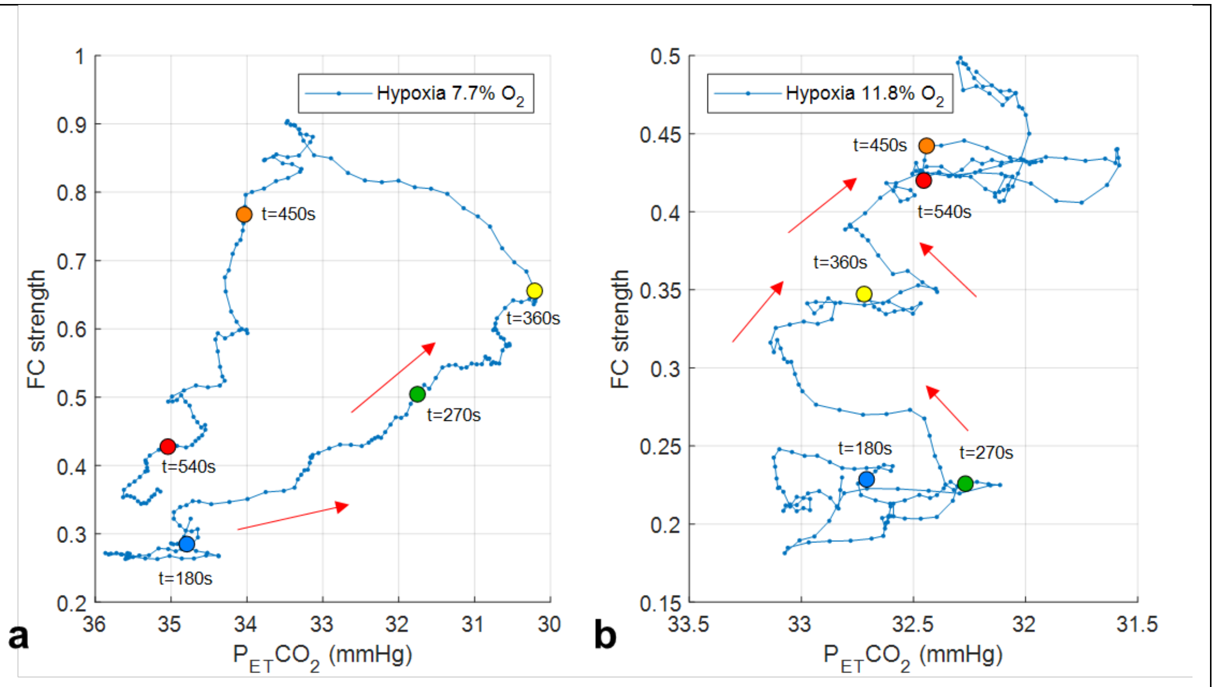


**Supplementary Figure 2. Hypoxia-induced FC changes demonstrate limited correspondence with $PetCO_2$**

**a,b**, Hypoxia-responsive FC strength, plotted as a function of $PetCO_2$ under severe ($FiO_2$ = 7.7%) and mild ($FiO_2$ = 11.8%) hypoxia, respectively.

Colored dots (blue to red) indicate five time points from 180s to 540s (90-second intervals), illustrating temporal progression alongside red arrows. In panel **a**, the blue and yellow dots mark the onset and offset of severe hypoxia, respectively. In panel **b**, all dots represent time points under hypoxia, as mild hypoxia was applied throughout. Baseline $PetCO_2$ levels prior to hypoxia were 35.0 ± 0.5 mmHg (averaged over 180 s) and 35.0 ± 0.3 mmHg (averaged over 240 s) for the severe and mild conditions, respectively. Compared to the pronounced changes in $PetO_2$, the fluctuations in $PetCO_2$ during this experiment were minimal.
